# Allelic polymorphism shapes community function in evolving *Pseudomonas aeruginosa* populations

**DOI:** 10.1101/644724

**Authors:** Sheyda Azimi, Aled E. L. Roberts, Shengyun Peng, Joshua S. Weitz, Alan McNally, Samuel P. Brown, Stephen P. Diggle

## Abstract

*Pseudomonas aeruginosa* is an opportunistic pathogen that chronically infects the lungs of individuals with cystic fibrosis (CF) by forming antibiotic resistant biofilms. Emergence of phenotypically diverse isolates within CF *P. aeruginosa* populations has previously been reported, however, the impact of heterogeneity on social behaviors and community function is poorly understood. Here we describe how this heterogeneity impacts on behavioral traits by evolving the strain PAO1 in biofilms grown in a synthetic sputum medium for 50 days. We measured social trait production and antibiotic tolerance and used a metagenomic approach to analyze and assess genomic changes over the duration of the evolution experiment. We found that (i) evolutionary trajectories were reproducible in independently evolving populations; (ii) over 60% of genomic diversity occurred within the first 10 days of selection. We then focused on quorum sensing (QS), a well-studied *P. aeruginosa* trait that is commonly mutated in strains isolated from CF lungs. We found that at the population level (i) evolution in sputum medium selected for decreased production of QS and QS-dependent traits; (ii) there was a significant correlation between *lasR* mutant frequency, the loss of protease and the 3O-C12-HSL signal, and an increase in resistance to clinically relevant β-lactam antibiotics, despite no previous antibiotic exposure. Overall, our findings provide insights into the effect of allelic polymorphism on community functions in diverse *P. aeruginosa* populations. Further, we demonstrate that *P. aeruginosa* population and evolutionary dynamics can impact on traits important for virulence and can lead to increased tolerance to β-lactam antibiotics.

## Introduction

The cystic fibrosis (CF) lung is a spatially complex and inflamed environment that provides beneficial growth conditions for a number of bacterial species, including the opportunistic pathogen *Pseudomonas aeruginosa* (1-5). Chronic infection of CF lungs with highly adapted and antibiotic resistant biofilms of *P. aeruginosa* is a major cause of lung function decline, which results in a concomitant increase in morbidity and mortality in individuals with CF (3, 4, 6-11). Longitudinal studies of chronic CF infections with *P. aeruginosa* have revealed that patients become infected at a young age with an environmental or transmissible isolate that evolves and adapts over time to the lung environment (12-18). Studies on *P. aeruginosa* populations isolated from individual lungs, have demonstrated divergent evolution, resulting in heterogeneous populations of *P. aeruginosa* within patients (16, 17, 19). This genetic adaptation and diversification is likely to impact on levels of pathogenicity and the efficacy of antibiotic treatment (16, 20), and could potentially impact how other species of microbe colonize the CF lung (4, 21-27).

Studies on explanted CF lungs have shown that the spatial structure found within lungs, and physical separation of infecting isolates, plays a role in generating the vast phenotypic and genotypic heterogeneity seen within *P. aeruginosa* populations in individual patients (16, 17, 28, 29). Major adaptations of *P. aeruginosa* to the CF lung include alginate production, loss of quorum sensing (QS), hypermutability and increased resistance to antimicrobials (7, 8, 14, 30, 31). Heterogeneity in *P. aeruginosa* populations has also been explained by early divergent evolution and adaptation to differential ecological niches (32) and recombination between isolates residing in the airways (16, 33), however, the exact mechanisms leading to heterogeneity have yet to be fully elucidated. Further, whilst it is accepted that genomic heterogeneity arises in CF chronic lung infections, it remains unknown how genotypic changes shape community functions within whole *P. aeruginosa* populations. Understanding what drives community structure and function remains a key goal in microbial ecology, because overall community function is determined by all the individuals in the population.

In this study, we hypothesized that the combination of wild-type and mutated alleles in populations shapes community functions, which could result in clinically-relevant outcomes such as increased or decreased antibiotic tolerance and traits important for virulence. To test this, we first evolved *P. aeruginosa* PAO1 in biofilms on plastic beads (34) for up to 50 days. This bead biofilm system has previously been successfully used to study genetic adaptation and phenotypic diversity of *Burkholderia cenocepacia* and *P. aeruginosa* to different environmental conditions (35). Our study differs from this previous work, in that our major focus was on community function within populations rather than on specific isolates and defining ecological niches. We grew our bead associated biofilms in a synthetic CF sputum media (SCFM), which recapitulates the chemical environment found in CF sputum (36-38). Our study therefore generated phenotypically and genotypically heterogeneous populations of *P. aeruginosa* in a spatially structured environment chemically relevant to CF sputum. We utilized these heterogenous evolved populations to study the changes in the functional community phenotypes instead of the traditional approach of working with single evolved isolates. We used a metagenomic approach to assess genetic alterations within evolving populations, and we monitored fluctuations in allele frequency during the selection process. To determine the impact of genomic heterogeneity within populations on various phenotypes, we assessed collective phenotypic traits (community function) of the evolved populations.

One of the most commonly described adaptations of *P. aeruginosa* to the CF lung is the loss of the *las* QS system, predominantly through point mutations, frameshifts and deletions in the *lasR* gene (17, 39-41). We used our evolved populations to specifically focus on the impact of *lasR* mutation frequency on QS phenotypes. The *lasR* gene encodes the LasR transcriptional regulator, which binds the QS signal *N*-(3-oxo-dodecanoyl)-L-homoserine lactone (3O-C12-HSL) (42-45). LasR-bound 3O-C12-HSL controls the transcription of approximately 10 % of the *P. aeruginosa* genome, including a number of genes involved in social behaviors, pathogenesis, antibiotic resistance and biofilm formation (46-49). Despite a number of previous studies which describe the changes and adaptation of various lineages of *P. aeruginosa* in CF lungs (14, 50-53), it remains unclear how polymorphisms in the *lasR* gene impacts on community function within evolving heterogenous *P. aeruginosa* populations. This is because most previous studies that focused on within host adaptation of *P. aeruginosa* used single colonies isolated from temporal CF sputum samples and not whole populations (12, 14, 15, 19, 54).

Overall we found that (i) evolutionary trajectories were reproducible between independently evolving populations and that over 60% of genomic changes in populations occurred within the first 10 days of selection; (ii) after 30 days of evolution in SCFM, the evolved communities displayed an increase in *lasR* mutant frequency and a decrease in QS-dependent traits; (iii) there was a significant correlation between *lasR* mutant frequency, the loss of social traits and an increase in tolerance to β-lactam antibiotics. Our findings provide insights into how allelic polymorphism and population heterogeneity in general, can impact on phenotypes and community functions within evolving *P. aeruginosa* populations. Further, we demonstrate that changes in *P. aeruginosa* population dynamics can alter factors associated with virulence and provide explanations for increased antibiotic tolerance, even in situations when antibiotics have not been used.

## Results

### Genomic variation in evolving biofilm populations over 50 days selection in SCFM

We evolved the *P. aeruginosa* strain PAO1 for 50 days (≈ 800 generations) in biofilms using a previously described biofilm bead method (34), and a growth medium that chemically mimics CF sputum (SCFM), and where the physiology of *P. aeruginosa* is similar to when grown in human sputum (36-38). Our experimental evolution approach contained four-independent replicate lines (Fig. S1). We collected and stored biofilm evolved populations after 10, 20, 30, 40 and 50 days of evolution (Rounds 1 to 5: R1 – R5). We used the Illumina MiSeq platform to deep sequence evolved populations in order to determine genomic changes through time. We also sequenced our laboratory PAO1 ancestral strain, and after *de novo* genome assembly of this strain, we mapped the sequence reads of the evolved populations to the ancestor in order to detect SNPs (55). Our SNP calling analysis, combined with an analysis of allele frequency, revealed that in all four independent replicate lines, an average of 282 ± 13 SNPs occurred in the populations after 10 days of selection (Fig. 1A). We found that around 60 % of these SNPs, were present through all other rounds of selection (Fig. 1B; Fig. S2). This suggests that the evolutionary trajectories of biofilm growth in SCFM are similar in independently evolving populations, and that the major genetic heterogeneity in evolving populations occurs during the early phases of selection.

### SNP frequency in genes involved in social traits fluctuates over time

In our evolution experiment, we found emergence of polymorphisms in 45 genes involved in various physiological functions (Fig. S3; Table S1). We found that between 10-25% of SNPs were fixed (frequency of 1) in the populations over 50 days of selection across all four replicate evolution lines (Fig. S4). When we focused on the allele frequency and not the number of positions altered in each coding region, we found that the frequency of SNPs in genes involved in different traits changed during the course of the experiment (Fig. 2; Fig. S3; Fig. S4 and Fig. S5). The genes highlighted in Fig. 2 are genes that have previously been shown to be commonly mutated in *P. aeruginosa* CF isolates (9, 15, 16, 29, 30, 56).

**Figure 1.**
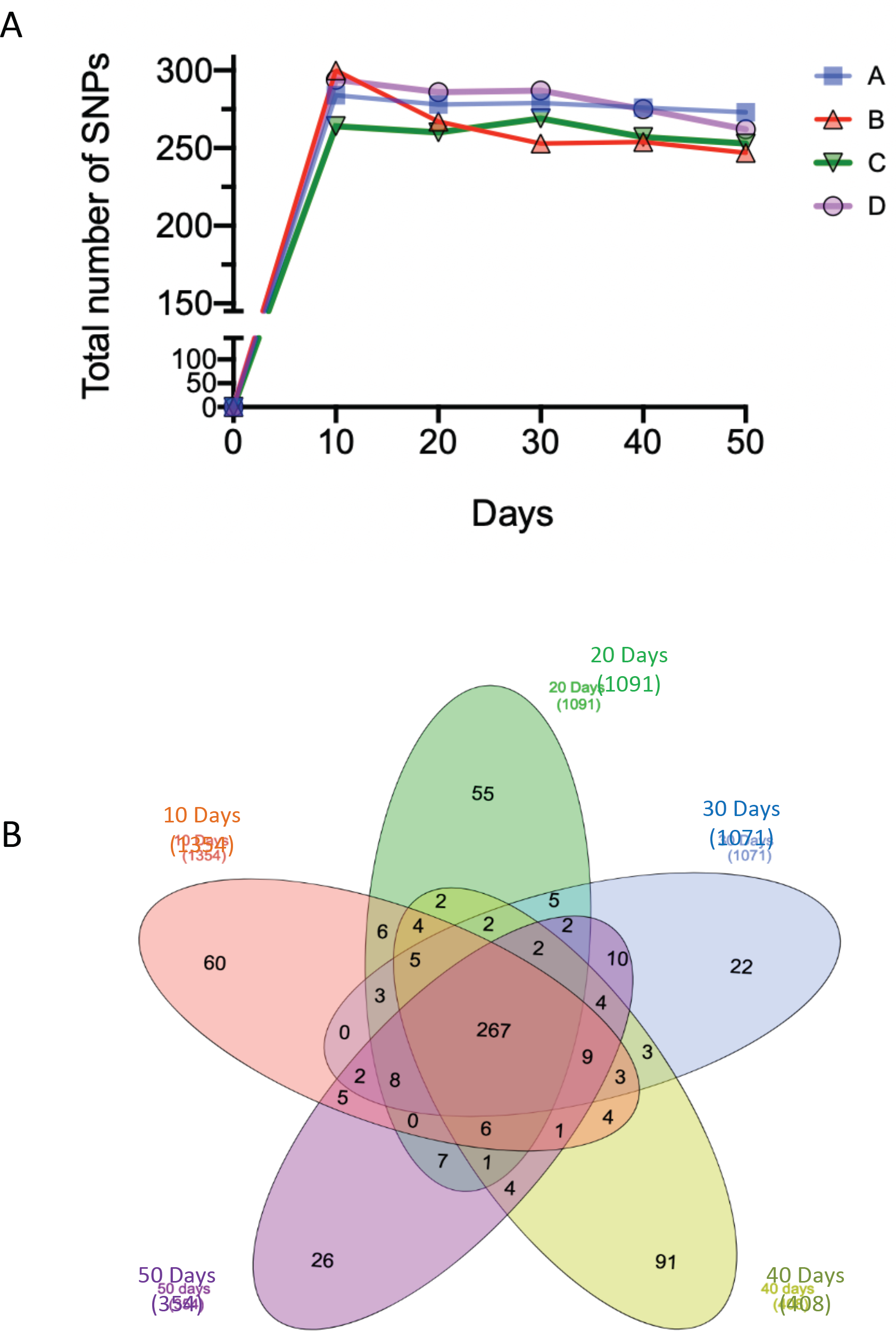
The evolutionary trajectory of *P. aeruginosa* during biofilm growth in SCFM. The evolution trajectories were similar in four independent replicate lines, and the major genetic heterogeneity in the evolved populations occurred during the first 10 days of selection. (A) SNP calling analysis showed that the four independently evolved populations had more than 80% similarity (267 shared SNPs) in genomic changes in the populations over the 50 days of selection. (B) The Venn diagram shows the shared SNPs between all 4 replicate lines, over 50 days of selection. Numbers in brackets, (i.e 1354 for 10 days), represent the combined number of SNPs in each of the four evolved lines. The majority of genetic variation within the populations occurred within the first 10 days of selection. Each circle represents the total number SNPs for each independently evolved line.

**Figure 2.**
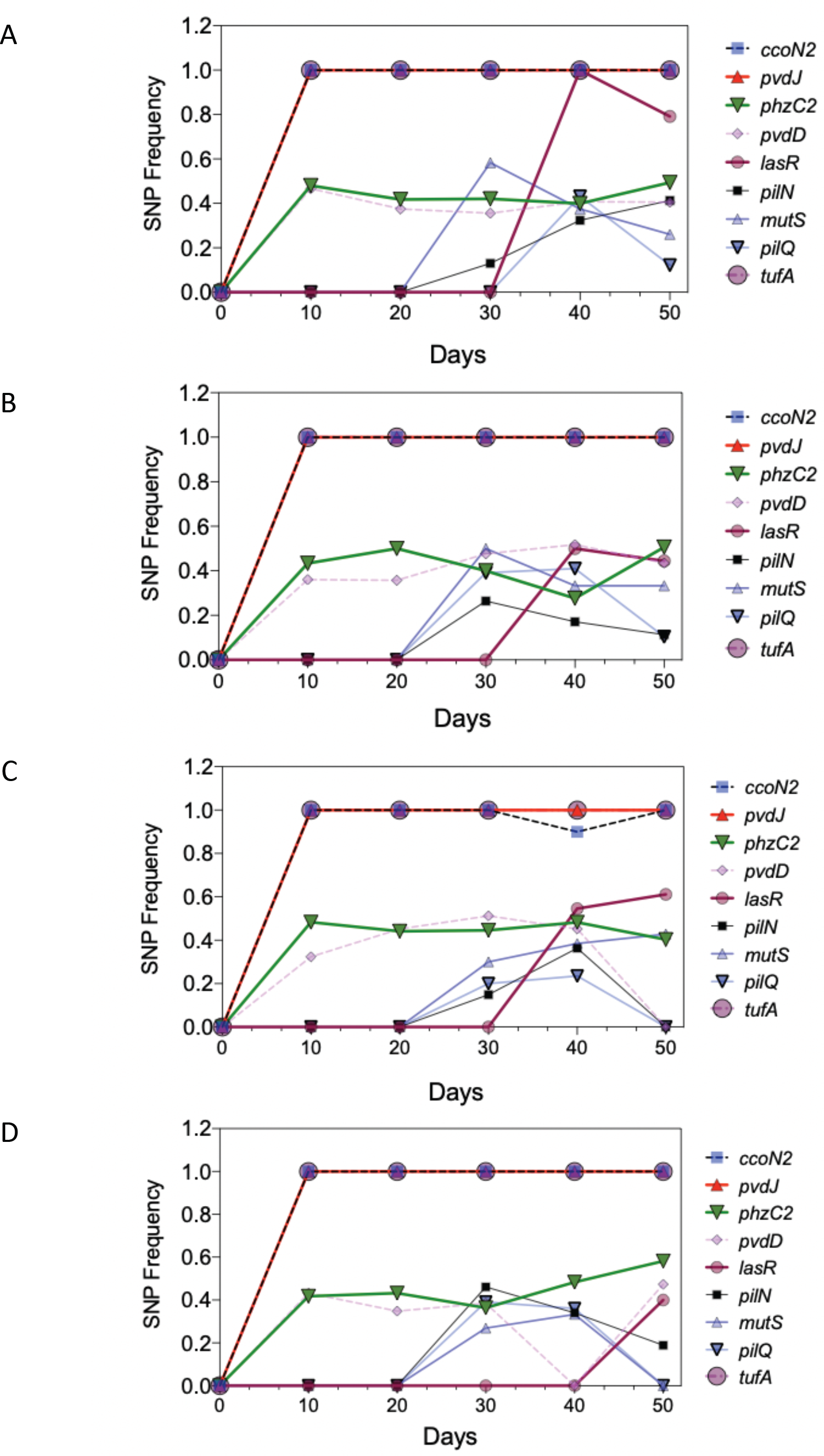
Allele frequency of SNPs changes over the course of selection. Emergence of nonsynonymous SNPs in genes involved in social traits (QS: *lasR*), oxidative respiration and DNA mismatch repair, motility and iron chelation occurred after 20 and 30 days of selection; and fixed SNPs in genes such as elongation factor *tufA*, Cytochrome c oxidase subunit (*ccoN2*) and pyoverdine biosynthesis protein *pvdJ* emerged after 10 days of selection and were fixed in the populations over 50 days (A-D represents replicate evolving lines).

The allele frequencies of nonsynonymous SNPs in *ccoN2* (PA1557), and synonymous SNPs in *pvdJ* (PA2400) and *tufA* (PA4265) became fixed in the population at a frequency of 1, while the frequency of nonsynonymous SNPs in *phzC2* (PA1901) and *pvdD* (PA2399) fluctuated between 0.4-0.5 in different rounds of selection. We detected a number of SNPs occurring in *mutS* (PA3620), *pilQ* (PA5040), *pilN* (PA5043) at 10, 20 and 30 days of selection in all four independent evolved lines. We observed an increase in *lasR* (PA1430) mutant allele frequency between 30-40 days of selection (Fig. 2; Fig. S3 and Fig. S5).

### Accumulation of SNPs shapes community functions in evolved *P. aeruginosa* populations

We next examined the production of phenotypic social traits in evolved populations in order to determine changes in community function of the genetically heterogenous evolving populations. We measured levels of biofilm formation, QS signals, total protease and the siderophores pyoverdine and pyochelin. We observed a small but significant increase in biofilm formation by evolved populations when compared to the PAO1 ancestor (Fig. 3A). After 30 days of selection, production of total protease (Fig. 3B) and the 3O-C12-HSL QS signal (Fig. 3C) decreased in evolved populations, however, the levels of C4-HSL signal (Fig. 3C), and pyochelin and pyoverdine (Fig. 3D) did not follow this trend, and any changes were generally not significantly different from the values of the ancestor strain. We also observed an increase in colony morphology types (morphotypes) in the evolved populations starting after 20 days of selection (Fig. S6).

**Figure 3.**
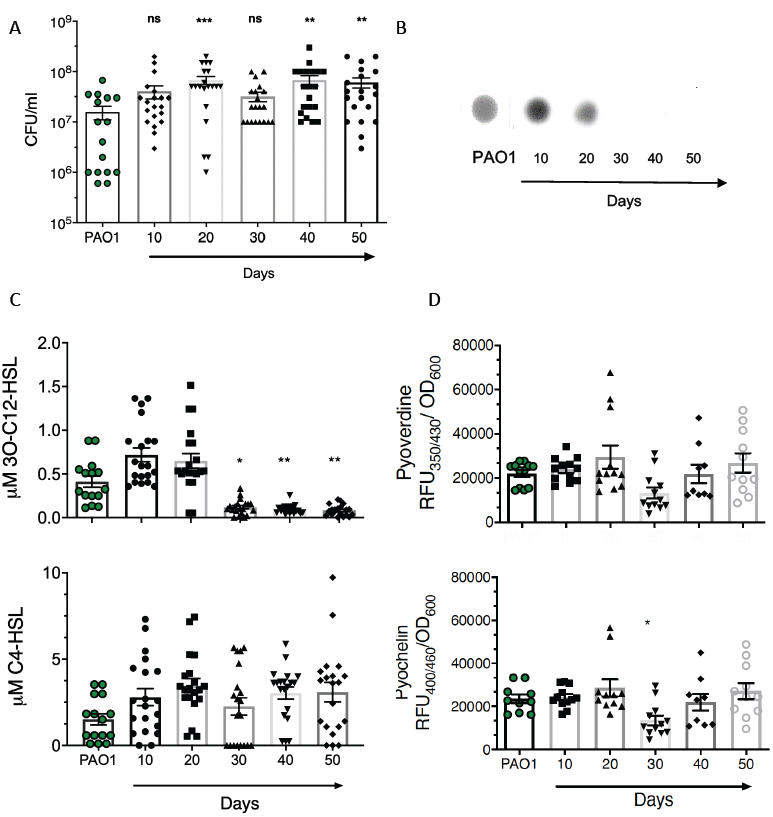
Phenotypes of evolved populations of PAO1. (A) Increase in biofilm formation by evolved populations of PAO1 (Kurskal-Wallis, Dunn’s multiple comparison to PAO1, *p* < 0.05, n=5. Errors bars indicate the standard error of the mean, SEM); (B) The total protease activity was reduced at 30 days of selection; (C) Production of 3O-C12-HSL significantly decreased at 30 days of selection (Kruskal-Wallis, uncorrected Dunn’s test multiple comparison to PAO1, *p* < 0.05, n=5. Errors bars indicate the SEM) and there were no significant changes in the levels of C4-HSL; (D) There were no significant changes in siderophore production in evolved populations compared to the PAO1 ancestor (Kruskal-Wallis, uncorrected Dunn’s test multiple comparison to PAO1, *p* > 0.05, n=5). Each dot represents the mean value of one independent experiment for each evolved population at each round of selection. All experiments were performed in 3 technical replicates.

To determine whether the emergence and increase in frequency of SNPs in the populations impact upon community function, we used a linear regression model. We focused on changes in the allele frequency detected in the QS regulator LasR. We chose *lasR* because the genes and phenotypes it regulates in *P. aeruginosa* are well understood (30, 40, 44, 47, 57) and also, *lasR* mutants are regularly isolated from CF sputum (13, 50, 58). We detected a non-synonymous SNP (V208G) in the DNA binding domain of LasR between 30-40 days of selection (Fig. S5). Interestingly, SNPs in the same position (V208) have been identified in *P. aeruginosa* isolates collected from CF sputum samples (40). V208 is adjacent to the D209 residue in the LasR DNA-binding domain (59, 60), suggesting an impact on the structure of LasR and its DNA binding affinity. We assessed the impact of *lasR* V208G SNP frequency on changes on two QS-dependent phenotypic traits; production of 3O-C12-HSL signal and total protease. We found a significant negative correlation between the frequency of the V208G *lasR* SNP in the whole-evolved populations and the total protease activity of evolved populations. We found that 87% (R^2^=0.8704, F=20.15; *p=*0.0206) of the decreased protease activity was correlated with the accumulation of the *lasR* mutation in the populations (Fig. 4A). We used the same analysis to determine whether *lasR* mutant accumulation impacts on 3O-C12-HSL production within populations. We found that only 53% of the decreased 3O-C12-HSL levels (R^2^=0.5363, F=3.469; *p=*0.1594) can be explained by the accumulation of *lasR* mutation in the populations (Fig. 4B). However, when we only included the last 30 days of selection in our analysis, the decrease in 3O-C12-HSL levels could be fully (100%) explained by the accumulation of *lasR* mutants (R^2^=1.0, *p=*0.0034) (Fig. 4B). Our analysis did not show any correlation between the frequency of the *lasR* SNP and changes in the production of C4-HSL (Fig. 4C) or biofilm formation (Fig. 4D).

**Figure 4.**
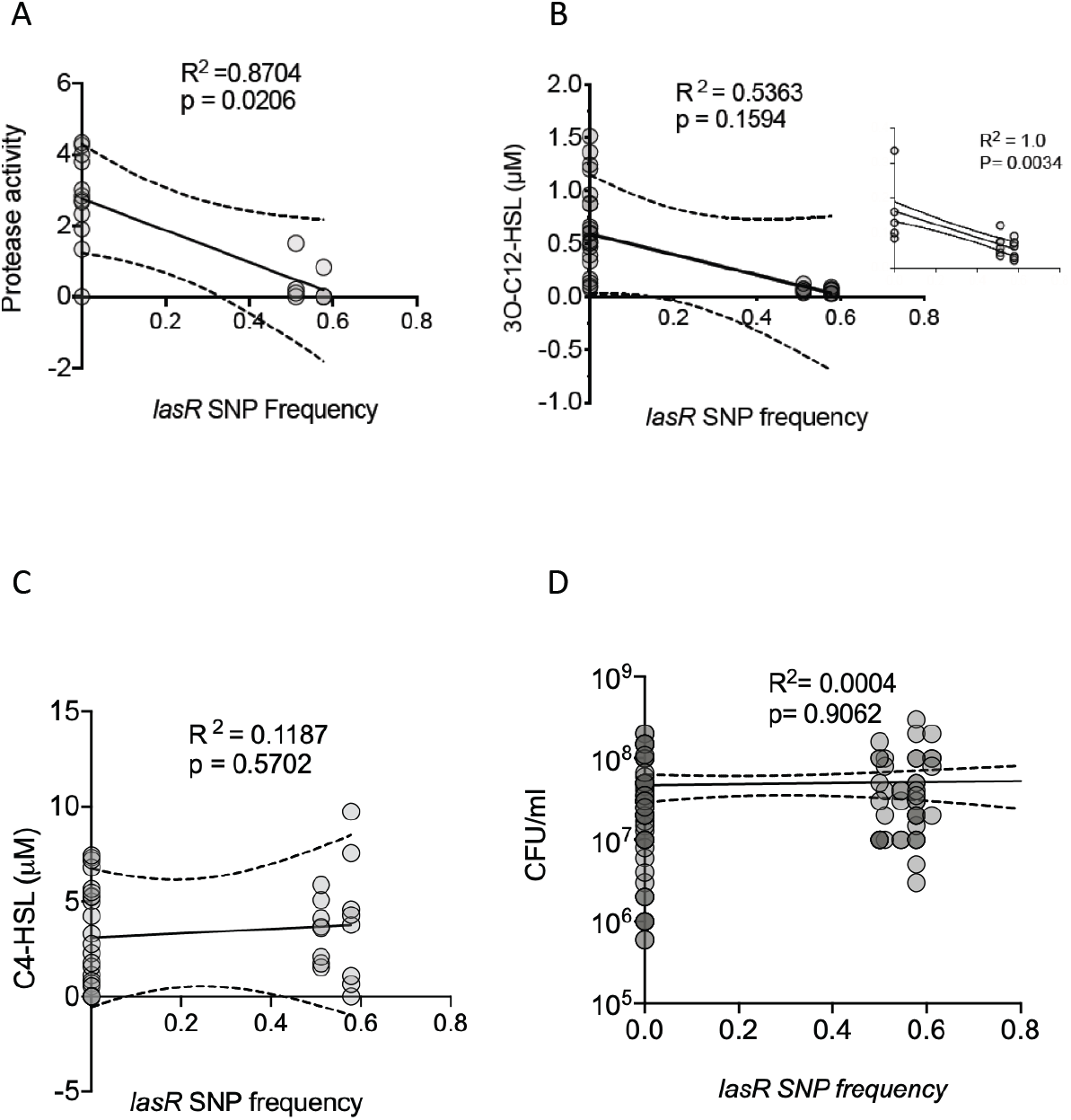
Loss of protease activity and a decrease in QS signal production in evolved populations can be explained by an accumulation of *lasR* SNPs. A linear regression model showed that the loss of total protease activity and the decrease in 3O-C12-HSL levels can be explained by the accumulation of *lasR* SNPs. (A) There is a significant negative correlation between *lasR* SNP frequency and total protease activity of the evolved populations (R^2^ = 0.7298, *p* = 0.04; n=8); (B) There is no significant correlation between 3O-C12-HSL levels and accumulation of the *lasR* SNP in the evolved populations (R^2^ = 0.5363, *p* = 0.1594; n=8). However, if only changes in 3O-C12-HSL levels after 30 days of selection is considered, there is a strong correlation between emergence and accumulation of *lasR* SNP in evolved populations (small inset: R^2^=1.0; *p =* 0.0034; n=8); (C) There are no significant correlations between accumulation of *lasR* SNPs in the evolved populations and levels of C4-HSL (R^2^ = 0.1187, *p* = 0.5702; n=8); (D) There is no significant correlation between *lasR* frequency and biofilm formation (R^2^ = 0.0004, *p* = 0.9062; n=8).

### Accumulation of *lasR* mutants in evolved populations leads to increased tolerance to β-lactam antibiotics

Previously it has been shown that *lasR* mutants display increased β-lactamase activity, and therefore increased resistance to β-lactam antibiotics (such as ceftazidime) (61) which are routinely used in CF clinics. To determine possible links between the loss of *lasR* function and changes in tolerance to routinely used antibiotics, we first assessed the antimicrobial susceptibility levels of the evolved populations. When we tested levels of antimicrobial susceptibility to six routinely used antibiotics for chronic CF lung infection, after 30 days of selection the evolved populations showed an increased tolerance (indicated by a decrease in the zone of inhibition) to three antibiotics; ceftazidime, piperacillin/tazobactam and meropenem which are all β-lactam class antibiotics (Fig. 5; Fig. S7). We then tested for correlations between frequencies of the *lasR* V208G SNP and resistance to ceftazidime and piperacillin/tazobactam. We observed a positive and significant correlation between accumulation of *lasR* mutants and the increased resistance to both ceftazidime and piperacillin/tazobactam (Fig. 5A and B). We also tested whether the increase in tolerance could be explained by an increase in biofilm formation. We found there was no significant correlation, suggesting that increases in biofilm formation do not necessarily translate to an increase in drug tolerance (Fig. S7C).

**Figure 5.**
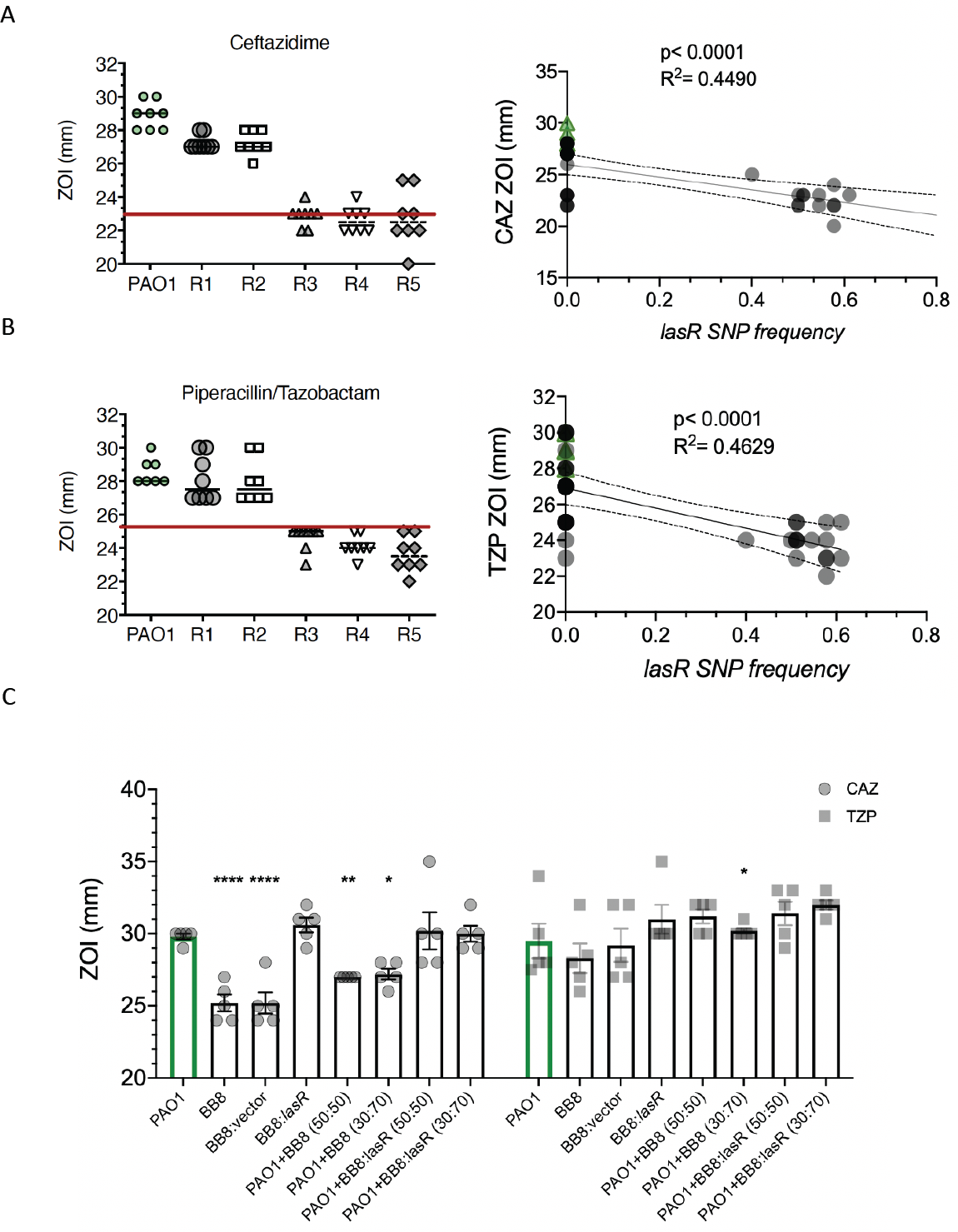
Increased resistance to ceftazidime and piperacillin/tazobactam in evolved populations is correlated with *lasR* SNP frequency. There is an increased resistance, indicated by a decrease in the zone of inhibition (ZOI) to ceftazidime (A) and piperacillin/tazobactam (B) in evolved populations after 30 days of selection (R3). The red lines represent the ZOI breakpoint (mm) for each antibiotic (BSAC guidelines). Below the lines indicates resistance. There is a positive correlation between increased resistance and the frequency of the *lasR* SNP in the evolved populations (n=8, Pearson correlation. Each dot represents the mean value for the zone of inhibition at each round of selection in all 8 independent experiments. (C) Complementation of a 50 day evolved isolate (BB8) containing a SNP in *lasR* with an intact *lasR in trans*, restores sensitivity to ceftazidime and piperacillin/tazobactam. Differential frequencies of B8 mixed with ancestral PAO1 significantly increased tolerance to ceftazidime (n=5, Two way ANOVA, multiple comparison to PAO1, Fisher’s LSD test, p <0.05. Errors bars indicate SEM).

To confirm that our evolved populations contained *lasR* mutants with increased β-lactam tolerance and that *lasR* frequency affects community function, we first selected and sequenced an evolved isolate (BB8) from the final round of selection (50 days) with no protease activity and which demonstrated increased antibiotic tolerance compared to the ancestor PAO1 (Fig. 5C). The sequencing of isolate BB8 revealed the same SNP in the *lasR* gene that we detected in the evolved populations (V208G) (Table S2). We complemented BB8 with an intact copy of the PAO1 *lasR* gene *in trans* on a plasmid and then assessed β-lactam tolerance levels. We observed a decrease in tolerance to ceftazidime and piperacillin/tazobactam in the *lasR* complemented BB8 isolate (Fig. 5C). Finally we tested whether changes in the starting frequency of *lasR* mutants in mixed populations impacted on community functions. We found that when BB8 was mixed with wild type cells, this resulted in an increased tolerance to ceftazidime (Fig. 5C) and a reduction in 3O-C12-HSL but not C4-HSL in mixed populations (Fig. S8).

## Discussion

Despite a number of recent studies focused on adaptive changes in evolved populations of *P. aeruginosa* in environments designed to mimic CF sputum (35, 62-66), there remain significant gaps in knowledge about how genomic and phenotypic diversity impacts upon on interactions within populations and on community-level phenotypes. In our current study, we focused on understanding how the evolution and coexistence of multiple lineages of *P. aeruginosa* shape community functions. To generate diversity, we performed a 50-day selection experiment, evolving PAO1 as biofilms in an artificial sputum medium (SCFM) (36-38). We observed (i) a rise in genetic diversity after 10 days of selection in biofilms grown in SCFM; (ii) up to 25% of SNPs became fixed in the population and carried in evolved populations through 50 days of selection; (iii) emergence and accumulation of SNPs in genes involved in motility, respiration, DNA mismatch repair, transcription and QS emerged between 10 and 40 days of selection and (iv) accumulation of *lasR* SNPs correlated with decreased protease activity, 3O-C12-HSL production and increased tolerance to β-lactam antibiotics despite no prior treatment with any antibiotic.

The divergent evolution of *P. aeruginosa* during chronic infection of CF lungs has been the focus of numerous studies on longitudinal collections of sputum samples. Analysis of single *P. aeruginosa* isolates from these collections has identified genomic signatures for adaptation to the CF lung environment (9, 14, 29, 50, 53, 67, 68). Similar studies on collections of *P. aeruginosa* isolates sourced from single sputum samples, have shown considerable phenotypic and genetic diversification of *P. aeruginosa* strains which group into clades (16, 17), and which may colonize different ecological niches in CF lungs (28, 69). All of these studies focused on the diversification, adaptation and heterogeneity of *P. aeruginosa* during chronic CF lung infection. However, the impact that this heterogeneity may have on collective phenotypes has been largely over-looked. We hypothesized that the combination of wild-type and mutated alleles in populations shapes collective phenotypes. We observed that the accumulation and frequency of various SNPs in heterogenous populations of *P. aeruginosa*, significantly impacts the collective phenotypes of evolved populations (Fig. 2; Fig. 3). Although we observed that the majority of genomic changes occurred during the first 10 days of selection, the functional outcome was only impactful after 20-30 days, where we observed an increase in biofilm formation and colony morphotypes and a significant decrease in total protease activity and production of 3O-C12-HSL by evolved populations.

The biofilm bead system has previously been used to examine the evolution of *P. aeruginosa* in biofilms (35), although there are significant differences between this and our study. First, the environmental conditions were different, with the previous study growing in a minimal media (M63) whilst we used SCFM. Secondly, we evolved PAO1 whereas the previous study evolved PA14 (35). Whilst both are popular model strains for *P. aeruginosa* study, they differ in a number of factors including virulence (5). Third, we focused primarily on the emergent properties of the whole community rather than individual isolates, which we show has important implications for signaling and antimicrobial tolerance. Finally, the previous study studied the emergence of *mutL* and *mutS* genes which may have contributed to the emergence of different individual morphotypes (35). We observed low frequency mutations in the *mutS* gene after 30 days (Fig. S3). As these were synonymous mutations, they did not appear to significantly increase mutation rate after 30 days of selection.

Our study also shows similarities with another recent study. Here the authors evolved PA14 in SCFM supplemented with mucin to create a structured environment (62). The main findings of this study was that mucin promoted diversification of *P. aeruginosa* between and not within populations. The authors found increases in tolerance to antibiotics in evolved populations and changes in growth, motility, pyocyanin production and biofilm formation. They did not present genomic evidence that underpinned these changes (62).

Our study identified genomic changes often observed in *P. aeruginosa* isolates taken from CF lungs including changes in secretion genes (*hcpA, hcpB, vgrG2a, vgrG4b*), motility genes (*pilQ* and *pilN*), iron aquisition genes (*pvdD, pvdJ*) and biofilm formation (*cdrA*). We also observed a consistent change in all lines and rounds of selection in anaerobic respiratory pathways (Fig. 2; Fig. S3). *ccoN2* (PA1557) encodes the cytochrome C oxidase subunit (ccb3 type). Increased expression of *ccoN2* in small colony variant has been previously noted during microaerophilic conditions (70, 71).

One of best-known signatures of *P. aeruginosa* adaptation to the CF lung environment, is mutation in the QS regulator LasR. Several studies on *P. aeruginosa* strains isolated from CF sputum samples, found significant genomic changes in *lasR*, including truncation, deletion, frame shifts and SNPs, with many resulting in a loss of function (30, 40, 41, 50, 72). Although there can be a high percentage of *lasR-*deficient isolates collected from patients, studies have also shown differential frequencies of functional and intact *lasR* alleles within patients (13, 73). Considering the importance of *lasR*-dependent social interactions for the fitness of *P. aeruginosa*, mutation-frequency of *lasR* within populations of *P. aeruginosa* could significantly impact on QS-dependent phenotypes and fitness at the population level (74-76). Previously we have demonstrated that an increase in the frequency of QS social cheats (*lasR* mutants) in defined populations of wild-type and *lasR* mutants, leads to a reduction in cooperation and virulence in mouse models of infection (74). These studies showed that even simple mixed genotype populations of *P. aeruginosa* can have a significant impact on community function, virulence and the outcome of infection.

In our current study, we observed a negative correlation between an increase in the frequency of the *lasR* V208G SNP in evolved populations and levels of protease activity and 3O-C12-HSL production (Fig. 4). Mutation of *lasR* in *P. aeruginosa* isolates from CF lungs has also previously been shown to be important for increased tolerance to β-lactam antibiotics such as ceftazidime (19, 56, 61). Here we examined whether the increase of *lasR* mutants in our evolved populations had an impact on levels of antimicrobial tolerance. Despite no prior treatments with antibiotics, we observed an increased tolerance to three antibiotics from the β-lactam family (Fig. 5). In contrast, we found no correlation between increased biofilm production by evolved populations and antibiotic resistance (Fig. S7). Resistance of *P. aeruginosa* to antibiotics in chronic infections such as CF or wounds, is often thought to be due to specific mechanisms such as efflux pumps or via production of excess polysaccharides such as alginate. Our findings suggest that the accumulation and frequency of genetic variants that might not traditionally be associated with resistance to drugs (e.g. QS mutants) within a heterogenous population, can alter phenotypes within populations that can result in important clinical repercussions.

Our work suggests that in the future, we should consider metagenomic and metaphenotypic assessments of *P. aeruginosa* populations collected from CF patients, rather than focusing on single colonies. This is because the phenotype of populations is dictated by the frequencies of various alleles in the populations. Focusing on just single isolates sourced from infections or long-term evolution experiments, results in particular strains being characterized with certain phenotypes, which misrepresents what is found in the population as a whole. It becomes particularly problematic in studies focusing on single colonies from longitudinal samples, and when genomic sequencing predicts how a strain genetically evolves over time during an infection. Our findings may also be particularly relevant when considering whether a *P. aeruginosa* infection is resistant or sensitive to antibiotic treatments. Our findings may extend to other infections caused by *P. aeruginosa* such as non-healing chronic wounds and they may also be relevant to other species of bacteria.

## Materials and methods

### Bacterial strains and growth conditions

For our experimental evolution, we used the PAO1 (University of Nottingham) strain of *P. aeruginosa.* For SCFM we followed the protocol provided in (36, 38). Briefly for the buffer base we prepared NaH_2_PO_4_ (1.3 mM), Na_2_HPO4 (1.2 mM), KNO_3_ (0.348 mM), K_2_SO_4_ (0.271 mM), NH_4_Cl (2.28 mM), KCl (14.9 mM), NaCl (51.8 mM) was prepared in 10mM of MOPS at pH=6.8, then the amino acids were added(36, 38). The Dextrose (3mM), L-lactic acid (9.3 mM), CaCl_2_*2 H_2_O (1.75 mM), MgCl_2_* 6H_2_O (0.606 mM) and Fe.SO_4_*7H_2_O (0.0036 mM) was added fresh every time the media was prepared.

### Long term experimental evolution

To assess how genomic diversity impacts on *P. aeruginosa* populations, we generated a diverse population using a long-term evolution experimental approach. We evolved the *P. aeruginosa* strain PAO1 in biofilms using plastic beads (34, 35, 77) (9×6 mm width) suspended in SCFM, in order to mimic a biofilm life cycle and a chemical environment similar to that found in CF lung sputum. To start the experimental evolution process, we first grew PAO1 on an LB agar plate. Then we inoculated a single colony of PAO1 into 3 ml of fresh SCFM (38), and incubated for up to 6 h to grow up to mid-log phase. We stored this mid-log phase cells as the ancestral PAO1 and compared all further phenotypic and genomic properties of the evolved populations to this. We inoculated mid-log phase cells to OD_600_ ≈ 0.05 into 4 tubes (in order to evolve separate independent lines named A-D) containing 3 ml of SCFM and a plastic bead. We incubated cultures for 24 h at 37 °C/200 rpm. After 24 h of incubation, we transferred bacterial covered beads into fresh tubes containing 3 ml of SCFM and a new bead. After each round the biofilms that formed on beads were composed of approximately 10^8^ cells. We then incubated again for 24 h at 37 °C/200 rpm. We continued the bead transfers for 50 days and stored biofilm portion of populations (attached to plastic beads) every 10 days (Rounds 1 to 5: R1-R5) (Fig. S1). We did not transfer or store the planktonic fractions at any point during the selection experiment.

### Deep sequencing of evolved populations

We extracted Genomic DNA from evolved populations after 18 h growth in SCFM; using DNeasy® Blood & Tissue Kit (QIAGEN) by following the manufacturer’s instructions. We prepared sequencing libraries using the NexteraXT protocol (Illumina), and sequenced in 24-plex on the Illumina MiSeq platform to obtain an approximate calculated level of coverage of 220-600× for each evolved population. A *de novo* assembly of the ancestral strain genome (*P. aeruginosa* PAO1 ancestor) was obtained using Spades with the –careful flag, and annotated using Prokka. We mapped reads of the evolved populations against the ancestral PAO1 genome using BWA(78). For SNP calling, the sequences were summarized using a MATLAB script for base quality, genomic position and mapping quality (55) script using bowtie2 (79), SAMtools (80) and BAMtools. To determine the allele frequency, we applied the breseq consensus model (81) to each of the samples collected. Later, for the variant sites, we used the composition in each nucleotide as the surrogacy for the allele frequency, and we excluded any allele frequency below 10% in each evolved population. Our deep sequence analysis was not designed to detect any insertions or deletions (INDELS), only SNPs. It is likely that INDELs were present in genes in our evolved populations.

### Measurement of biofilms formed on beads

To determine the levels of biofilm formation by each evolved population, we grew biofilms on plastic beads as previously described (34, 77). For each set of biofilm assays, we directly inoculated a 10 μl loop of frozen evolved populations (approximately 10^6^ cells) into 3 ml of SCFM and incubated at 37°C/200 rpm for 16 h. Then we measured the OD_600_, and diluted in 3ml SCFM to OD_600_ ≈ 0.05. Then 3 plastic beads were added to each tube. After 24 h growth at 37°C/200 rpm and biofilm formation. The beads were then washed 3× with 10 ml of PBS to remove any residual planktonic cells not bound to the plastic beads. Then we transferred each plastic bead into 1ml of PBS and sonicated the beads for 10 mins, using a bath sonicator to detach biofilm forming cells from the beads. We then serially diluted the cells and plated out onto LB agar plates for colony forming unit (CFU) calculations. To assess other phenotypic traits, a cell free supernatant of biofilm forming cells was prepared from liquid part of the cultures and corrected values for OD_600_ ≈1.

### Preparation of cell free supernatants

To assess the levels of protease, QS signals and siderophore production during biofilm formation, we collected 3 ml of the SCFM used in the biofilm assays. We use the media surrounding the bead biofilms; and measured the OD_600_ and adjusted it to 1 for all the cultures with SCFM. We then filtered the supernatants using 0.22 µm filters and used these cell free supernatants to measure phenotypic traits.

### Total protease activity

To assess the total protease activity of evolved populations, we used skimmed milk agar plates. We inoculated 10 µl of cell free supernatant from each evolved population onto skimmed milk agar plates (1.2% Bacto Agar, 0.015% of skimmed milk) alongside 10μl of 10μg/ml of proteinase K and supernatant of PAO1 as controls. The zone of clearance was scored based on appearance and measured with a ruler (in mm). We then imaged each plate using an Epson scanner at 800dpi. We then compared it to the zone of clearance produced by the PAO1 ancestor (82).

### Siderophore production

To measure the levels of the two main siderophores produced by *P. aeruginosa*, we used the cell free supernatants. 100 μl aliquots of cell free supernatant from evolved populations and the PAO1 ancestor was transferred into a black clear bottom 96 well plate (Corning). We measured the emission as Relative Fluorescent Units (RFU) using a Multi microplate reader (Tecan Infinite® M200 Pro). We measured the wavelengths at excitation of 400 nm/emission460 nm for Pyoverdine and 350/430 nm for Pyochelin (30, 83, 84). We corrected the values for pyoverdine and pyochelin to the absorption at (OD_600_) of the original cultures.

### Measurement of C4-HSL and 3O-C12-HSL produced by evolved populations

The cell free supernatants were used to determine the concentration of QS signals. We used two *E. coli* bioreporter strains to measure production of the two main signal molecules by the evolved population. The *E. coli* reporters pSB536 and pSB1142 were used to detect C4-HSL and 3O-C12-HSL respectively (85). We calculated signal levels based on standard curves fitted to the concentrations of synthetic 3O-C12-HSL and C4-HSL standards (Sigma) (86, 87).

### Mixed isolate experiments

For mixed constructed populations, we grew PAO1, the BB8 evolved isolate and BB8 complemented with *lasR* (BB8:*lasR*) and empty vector (BB8:vector) in SCFM to OD_600_ ≈ 0.5. We then mixed PAO1 and BB8 at two different starting frequencies (50:50 and 30:70) in 3 ml of SCFM. The cultures were then incubated for 16 h at 37°C/200 rpm. We measured and adjusted the OD_600_ of each culture to OD_600_ ≈ 1 and prepared a cell free supernatant for QS signal activity measurements.

### Antibiotic susceptibility assay

To determine the antibiotic susceptibility in evolved populations, we followed the British Society for Antimicrobial Chemotherapy (BSAC) guidelines (Version.14, 05-01-2015) using Isosensitest agar plates (Oxoid). We tested the susceptibility of evolved populations, PAO1 ancestral strain and the NCTC (10662) PAO1 strain to Gentamicin (10 μg), Meropenem (10 μg), Ciprofloxacin (1 μg), Ceftazidime (30 μg), Piperacillin/Tazobactam (85 μg) and Amikacin (30 μg) (Oxoid). The zone of inhibition and clearance in this method was compared to the available zone of inhibition breakpoints for susceptibility (mm) for each tested antibiotic based on BSAC guidelines.

### Determining colony morphology diversity in evolved populations

To determine the diversity in colony morphology in the biofilm evolved population, we used a Congo Red based agar media (1% agar, 1xM63 salts (3g monobasic KHPO4, 7g K2PO4, 2g NH4.2SO4, pH adjusted to 7.4), 2.5mM magnesium chloride, 0.4 mM calcium chloride, 0.1% casamino acids, 0.1% yeast extracts, 40 mg/L Congo red solution, 100 μM ferrous ammonium sulphate and 0.4% glycerol) (82). We recovered the evolved populations from beads and serially diluted the populations and then inoculated onto CRA plates alongside the PAO1 ancestor. We incubated the plates overnight at 37°C, and for a further 4 days at 22°C. The colonies were imaged using an Epson scanner at 800dpi.

### Complementation of *lasR*

To complement an evolved isolate (BB8) containing a *lasR* mutation with a functional *lasR* allele, we amplified a 920 bp product comprising *lasR* and 200 bp upstream of the *lasR* start codon that includes its native promoter, from genomic DNA isolated from wildtype ancestral PAO1. We cloned this PCR product into the shuttle vector pME6032, which replicates in both *E. coli* and *P.aeruginosa*, (88) by Gibson assembly (89) using the commercially available NEBuilder HiFi DNA Assembly Cloning Kit (New England Biolabs, Ipswich, MA). We introduced the *lasR* complementation construct into BB8 by electroporation (90) and plating on selective media containing 300 ug/mL tetracycline. We then tested susceptibility of the *lasR* complemented strains to ceftazidime and piperacillin/tazobactam using the BSAC method. For mixed constructed populations, we grew PAO1, the BB8 evolved isolate and BB8 complemented with *lasR* (BB8:*lasR*) and empty vector (BB8:vector) in SCFM to OD_600_ ≈ 0.5. We then mixed PAO1 and BB8 at two different starting frequencies (50:50 and 30:70) in 1 ml of PBS and tested for antibiotic tolerance by following the British Society for Antimicrobial Chemotherapy (BSAC) guidelines (Version.14, 05-01-2015) using Isosensitest agar plates (Oxoid).

### Statistical analysis

For statistical analysis of the phenotypic assays, we used GraphPad Prism 8.0. For analysis of SNP frequency, we used R package 3.6. We used the Interactive Venn (91) to analyze shared SNPs within and between evolved populations.

### Publication of genome sequencing

All sequences described in this manuscript have been uploaded to the NCBI SRA database (accession number PRJNA613708).

## Acknowledgements

For funding we thank the Human Frontier Science Program (RGY0081/2012) and Georgia Institute of Technology; The Cystic Fibrosis Foundation (DIGGLE18I0) to SPD; Cystic Fibrosis Foundation for a Fellowship to SA (AZIMI18F0); CF@latna for a Fellowship to SA (3206AXB). The National Heart Lung Blood Institute (R56HL142857) and The Simons Foundation (396001) to SPB. We acknowledge Jacob Thomas for help with *lasR* complementation and Freya Harrison and James Gurney for helpful comments on the work. We also thank three anonymous referees for their helpful suggestions for improving this manuscript.

## Supplementary figure legends

**Figure S1.**
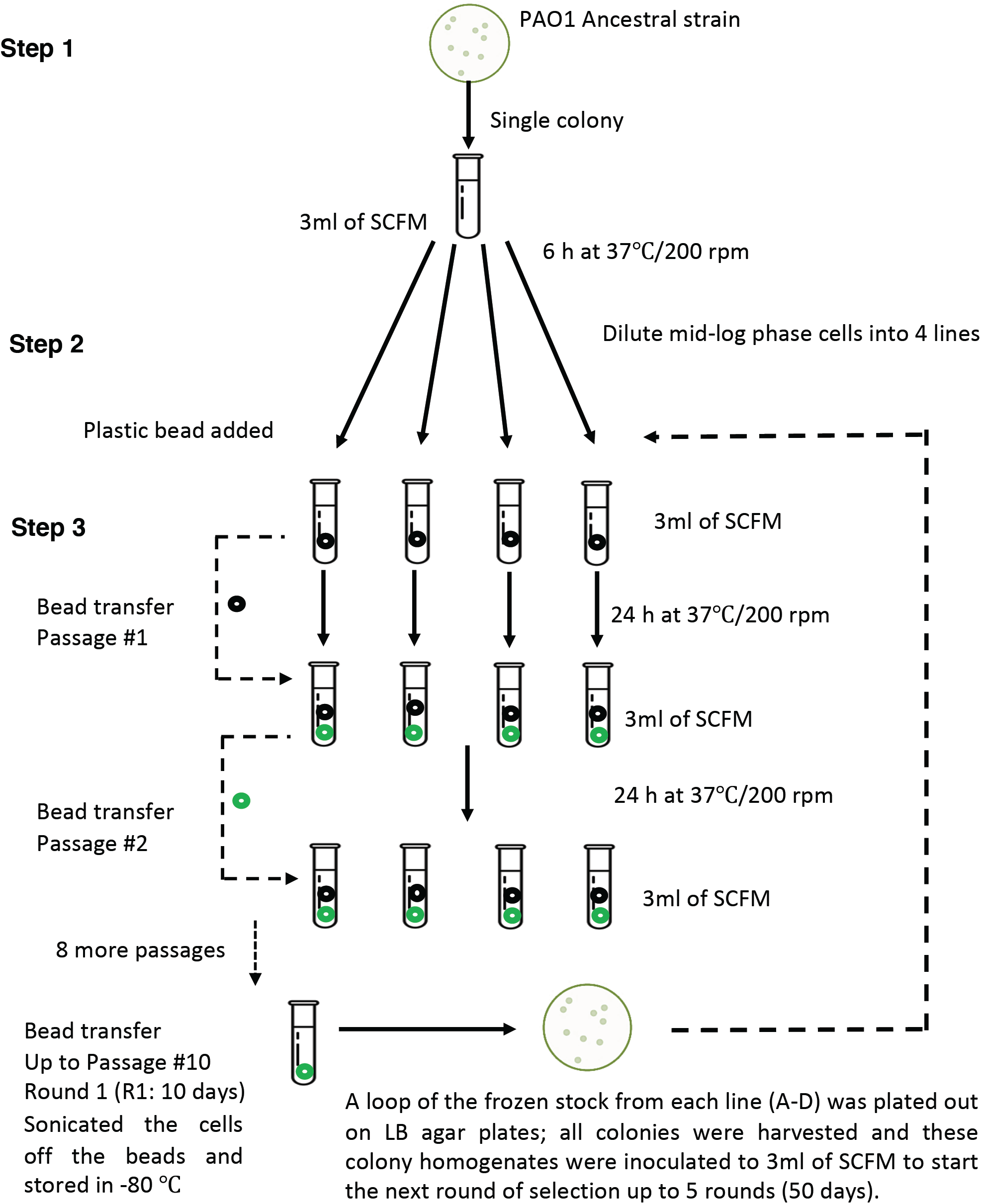
Long term evolution of PAO1 in biofilms and SCFM. PAO1 was grown in SCFM and inoculated into four independent replicate lines in the presence of one plastic bead (6×7 mm) to enable the bacteria to form biofilms on the surface. Every 24 hours the beads were transferred to a new tube with fresh SCFM media and a new bead. After 10 days, the bacterial populations were sonicated off the beads and stored as round 1 (R1). To start the next 10 days of selection, a 10 µl loop of bacteria (approximately 10^6^ cells) was used. This whole process was repeated 5 times to give 50 days (rounds 1 to 5) of selection.

**Figure S2.**
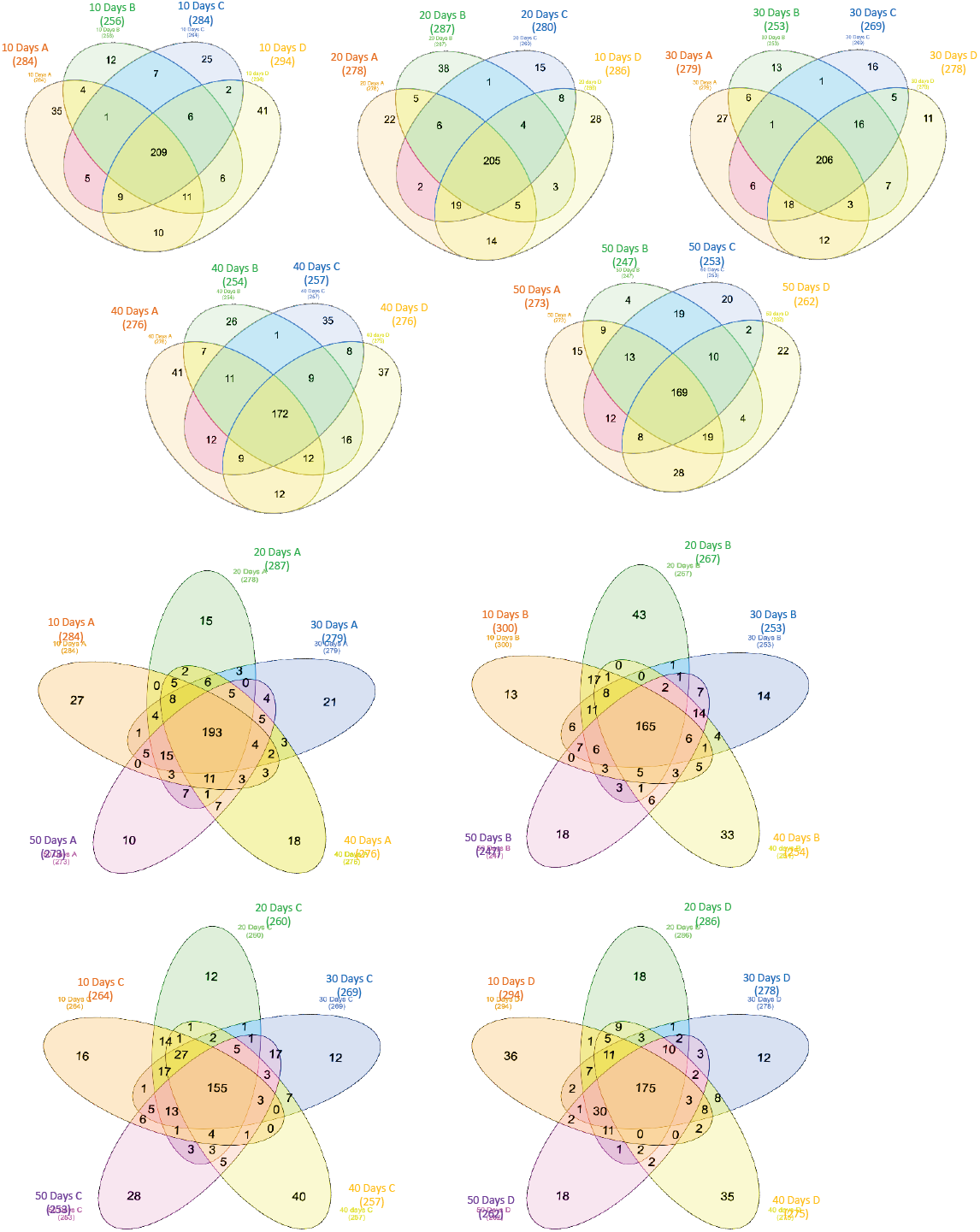
Number of SNPs shared in all evolved lines during 50 days of evolution in SCFM. We tested the number and percentages of the SNPS that occurred and are shared between various evolution lines at different rounds of selections. There are 207 SNPs that are shared between all 4 lines of selection (A-D) at day 10, 205 on day 20, 206 on day 30, 172 on day 40 and 169 on day 50 (upper panels). We observed that the majority of SNPs occurred during the first 10 days of selection and are shared and carried over to subsequent rounds of selection in all evolution lines.

**Figure S3.**
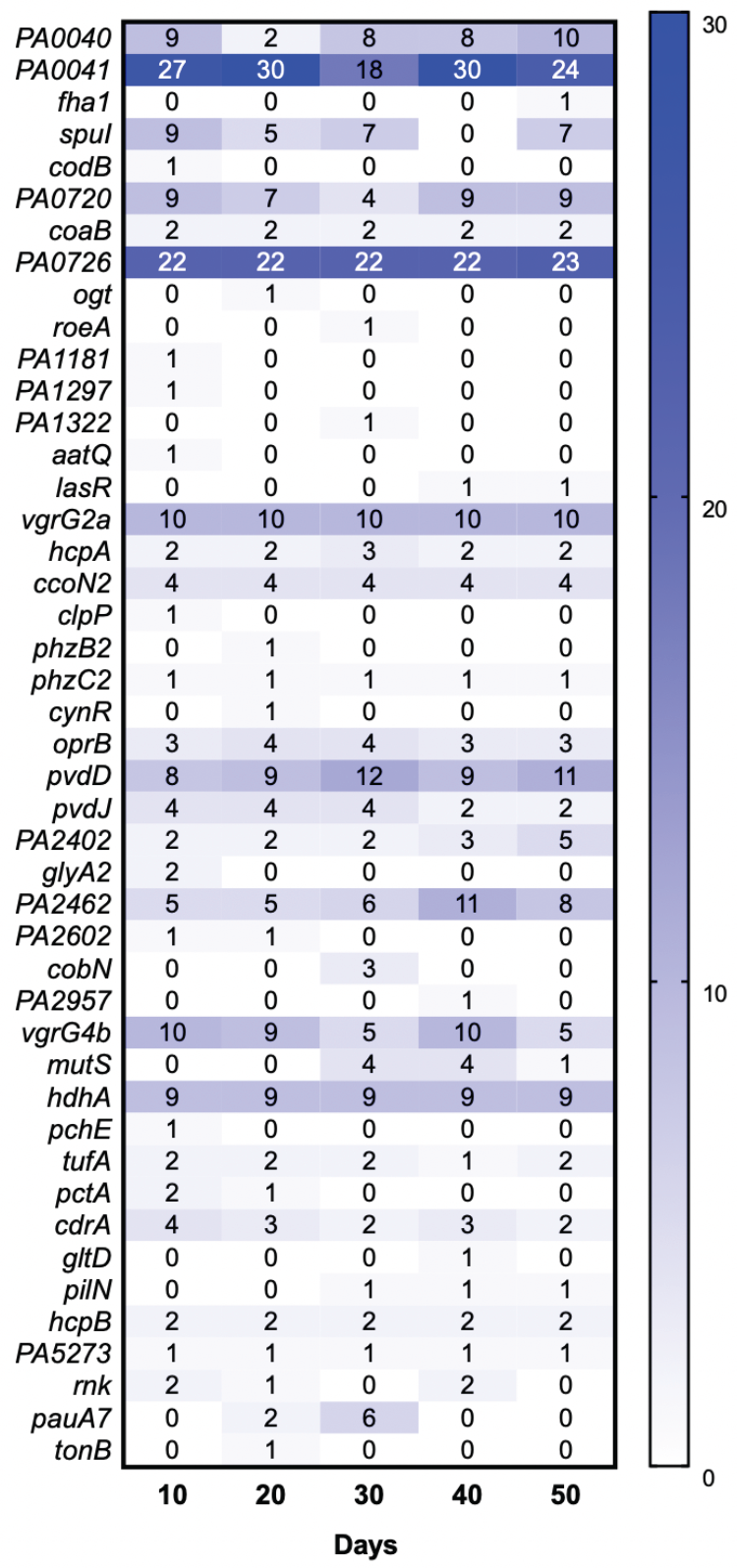
Number of SNPs in genes over 50 days of selection in SCFM. The emergence of polymorphisms in 45 genes involved in various physiological functions (Evolved line A shown as a representative for all evolved lines). The shaded boxes represent the abundance of SNPs in each gene.

**Figure S4.**
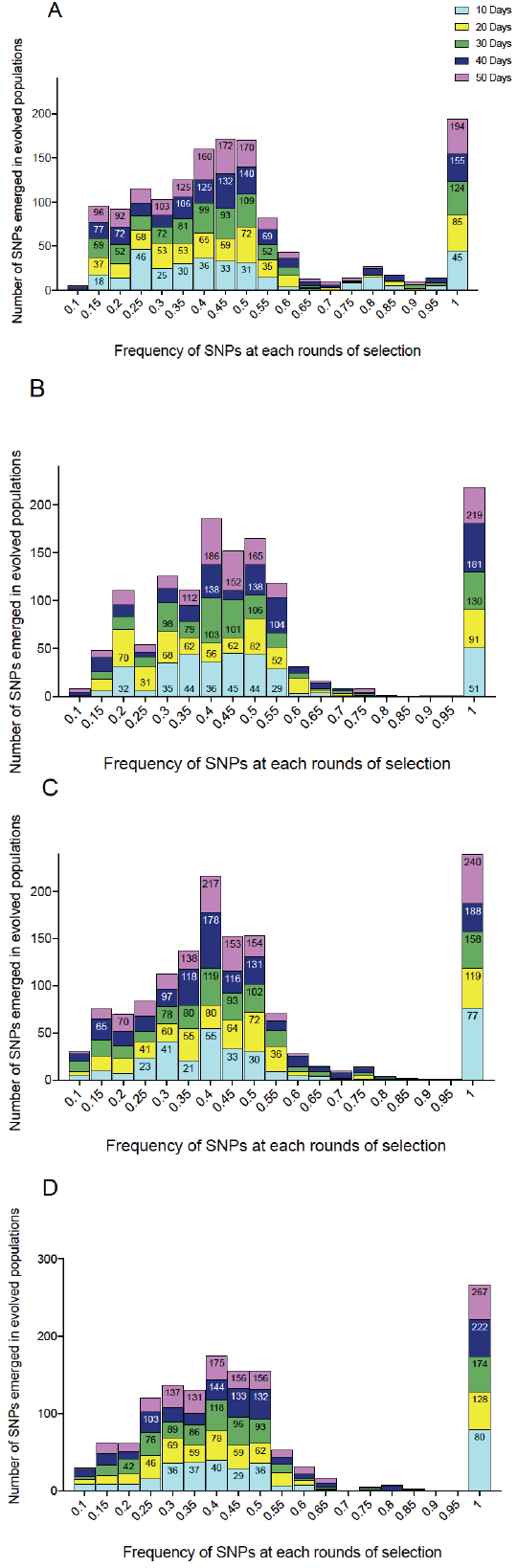
Abundance of differential allele frequencies in each evolved population over five rounds of selection. Different SNPS emerged after 10 days of selection at varying frequencies within the populations. Each round of selection contained varying frequencies of SNPs. Over 10-25% of SNPs were fixed in populations after 10 days of selection, across all four independently evolving populations (A-D).

**Figure S5.**
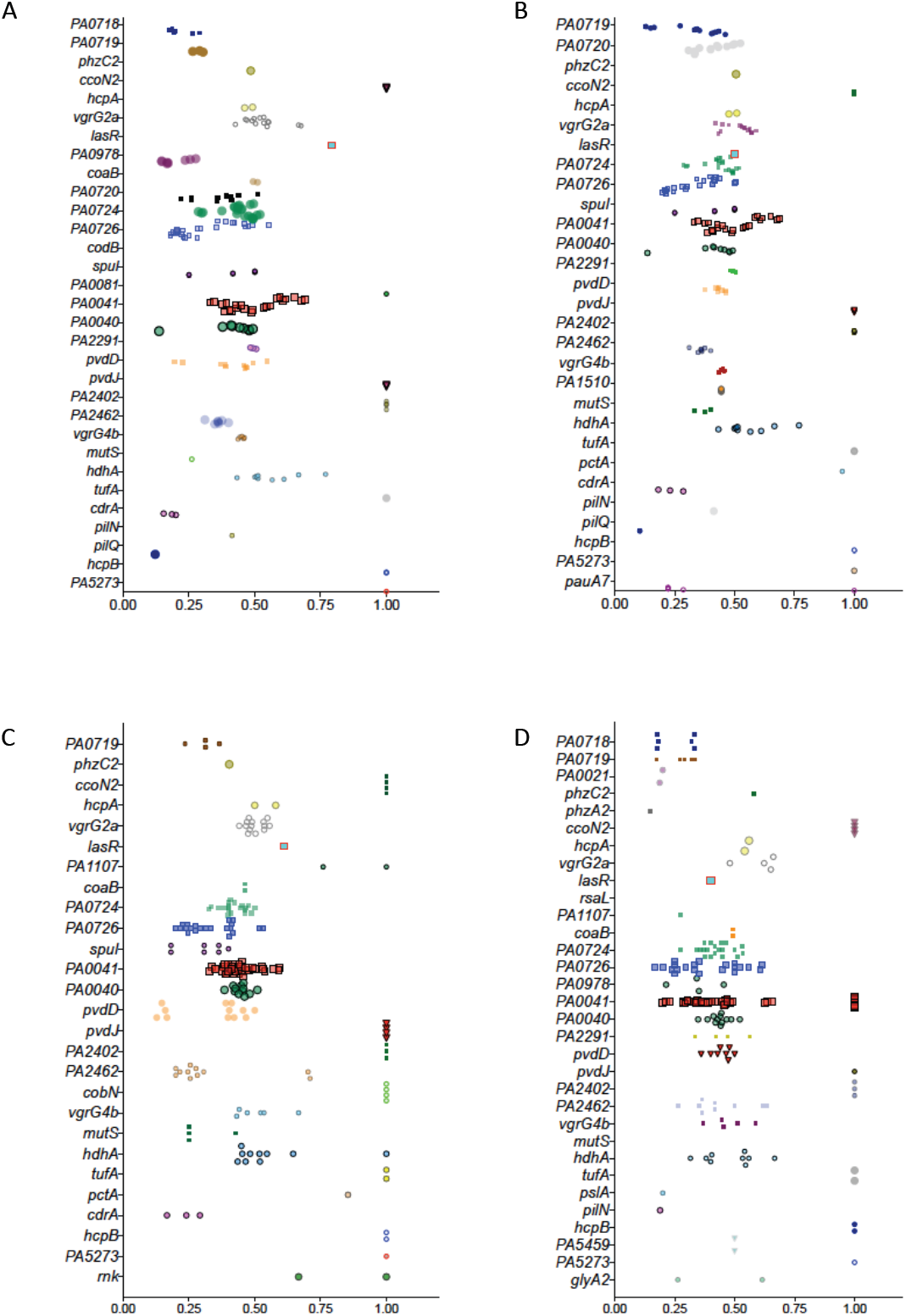
Number of SNPs in various genes after 50 days of selection. We found high levels of similarity in number and frequency of SNPs in all independent evolved lines after 50 days of selection. Each data point is representative of SNPs in each gene, at various frequencies in all four evolved lines.

**Figure S6.**
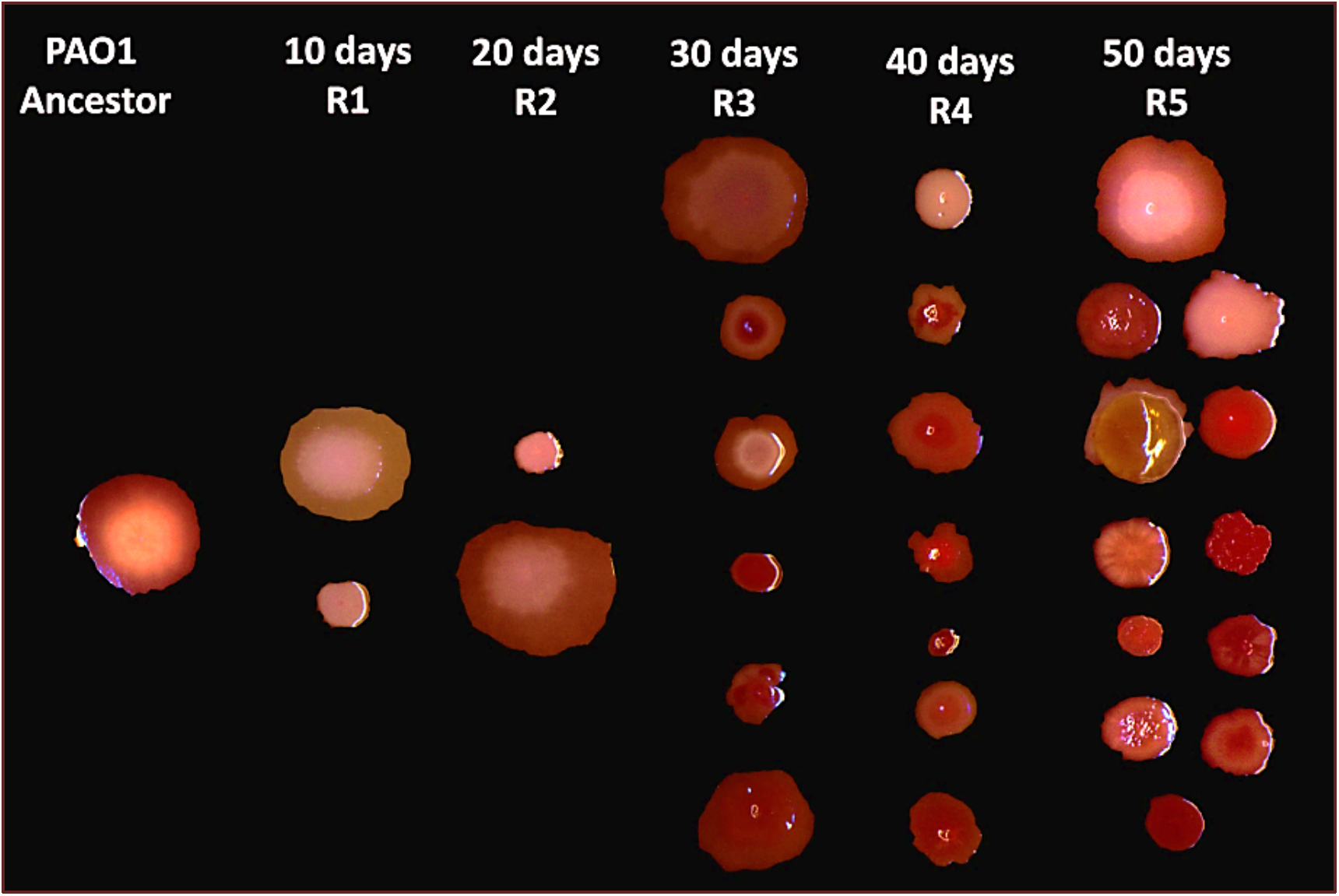
Increase in diversity of colony morphologies in evolving populations over time. Diversity of colony morphology increased over time in evolutionary lines. We used Congo Red Agar plates to highlight differences in morphologies between haplotypes.

**Figure S7.**
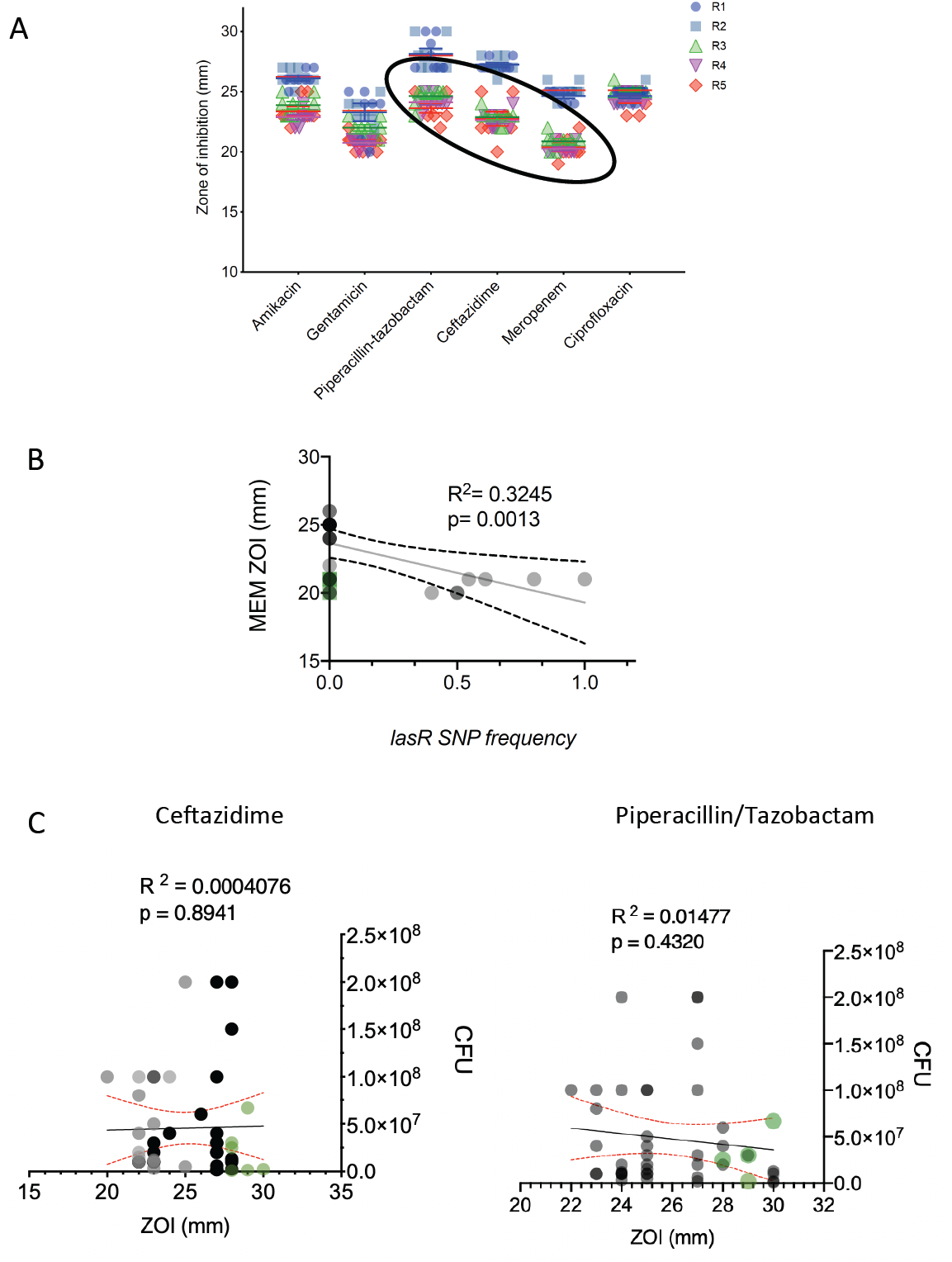
Increased resistance to b-lactam antibiotics in evolved populations. (A) There is a decrease in the ZOI to all b-lactam antibiotics after 30 days of selection in evolved populations. (B) There is an increase in resistance to meropenem after 30 days of selection in the evolved population, which significantly correlates to accumulation of the *lasR* SNP in the populations. However, the increased resistance does not pass the threshold to be considered clinically resistant to meropenem. (C) There is no correlation between the levels of biofilm formation or antibiotic resistance at the population level.

**Figure S8.**
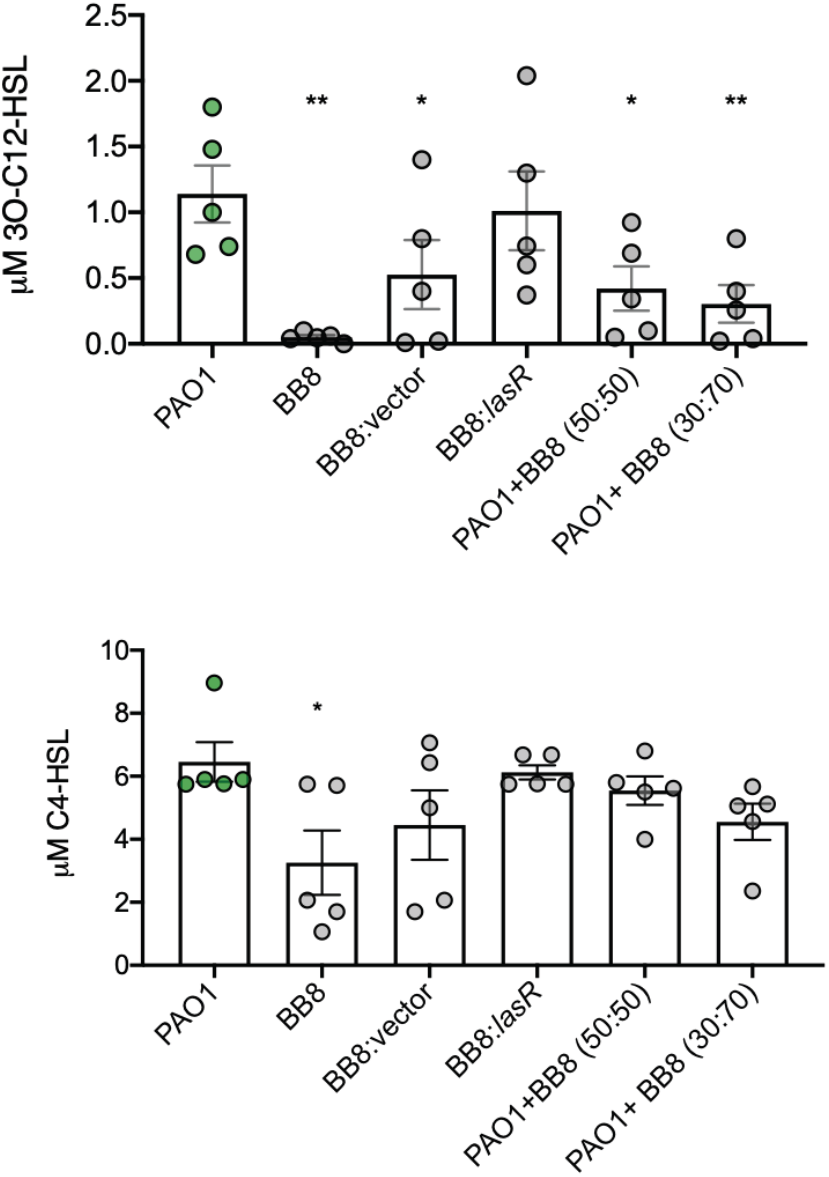
Increase in *lasR* mutant frequency changes the functional phenotype of synthetic mixed populations. Varying frequencies of BB8 isolate (V208G *lasR* allele) with the PAO1 ancestor, significantly alters the levels of 3O-C12-HSL and has no impact C4-HSL levels (Multiple comparison, One Way ANOVA, n=5).

**Table S1.** SNP positions and altered allele frequencies in four evolved lines over 5 rounds of selection.

**Table S2.** SNPs in the 50 day evolved isolate BB8.

